# Temporal Multi-Omic Analysis Uncovers Sex-Biased Molecular Programs Underlying Skeletal Muscle Adaptation to Endurance Training

**DOI:** 10.64898/2025.12.26.696612

**Authors:** Gina M Many, Christopher Jin, Nicholas Day, Gayatri Iyer, Gergory Smith, James Sanford, Akshay Bareja, David Jimenez-Morales, Kayleigh Voos, Damon Leach, Tyler Sagendorf, Xiaolu Li, Matthew Gaffrey, Isaac Attah, Hugh Mitchell, Mark Viggars, David Gaul, Kim M Huffman, Fácundo Férnandez, Michael P Snyder, Eric Ortlund, Wendy Kohrt, Matthew T Wheeler, William E Kraus, Karyn A. Esser, Bret Goodpaster, Charles F Burant, Christopher B Newgard, Andrea Hevener, Sue C Bodine, Wei-Jun Qian, Simon Schenk, Joshua Adkins, Malene E Lindholm, the MoTrPAC study group

## Abstract

**Background:** Exercise training is known to benefit health and reduce disease risk. While skeletal muscle adaptations are fundamental to many of the health benefits of exercise training, the common and sex-specific molecular regulators that mediate these adaptations remain to be fully elucidated.

**Methods:** To this end, we leveraged skeletal muscle multi-omics data generated by the Molecular Transducers of Physical Activity Consortium (MoTrPAC), where 6 month-old male and female rats endurance trained for 1, 2, 4, or 8 weeks. Our objective was to identify shared and sex-specific multi-omic molecular responses to endurance training in skeletal muscle, and relate them to phenotypic adaptations.

**Results:** We identified largely sexually-conserved transcriptomic and proteomic pathway enrichments in the *gastrocnemius*, which correlated with skeletal muscle responses from a published exercise study in humans. We uncovered sex-consistent post-translational modifications, including decreased oxidation of MYH2 and deacetylation of the β-oxidation enzyme HADHA. Pathway enrichment analyses revealed sex-specific remodeling across the acetylome, redox proteome, and phosphoproteome; females decreased mitochondrial protein cysteine oxidation and increased mitochondrial cristae proteins, indicative of enhanced redox buffering and mitochondrial efficiency. Despite decreases in cysteine oxidation of key mitochondrial proteins, females displayed increases in the cysteine oxidation of proteins involved in glucose catabolism relative to males after 8 weeks of training, suggestive of sex-biased subcellular reactive oxygen species generation. Males demonstrated earlier induction of mitochondrial transcripts and predicted activation of mTOR. Although the increase in mitochondrial protein abundance was more modest in males, there was greater oxidation of mitochondrial proteins in response to training compared to females.

**Conclusions:** This work shows a large portion of the adaptive response to endurance training in skeletal muscle is shared between females and males, while there are distinct and nuanced sex-specific adaptations that are evident, particularly at the level of post-translational regulation.

## Introduction

Endurance exercise training has long been a cornerstone of maintaining or improving whole-body health and reducing disease and mortality risk. While exercise training induces many physiological adaptations throughout the body^1,2^, adaptations specific to the skeletal muscle engaged during exercise (e.g. increased oxidative capacity) are fundamental contributors to the salutary effects of training. Epidemiological evidence shows that the association between being physically active and all-cause mortality differs between men and women, such that women have greater reductions from equivalent doses of leisure-time physical activity^3^. Historically, exercise studies have primarily included males, emphasizing the need to investigate potential differences in molecular adaptation between men and women.

Over the past six decades, concerted effort has gone into understanding what changes occur in skeletal muscle with exercise training, along with how these changes are regulated^4,5^. While this focus was initially on physiological and morphological changes in skeletal muscle, over the past ∼20 years, high-throughput, omics-based studies have sought to identify molecular signatures that drive the adaptive response. To date the most prevalent omics approaches used have assessed the epigenome (methylome), transcriptome or global proteome, with the most commonly identified adaptations being the ability of endurance training to improve mitochondrial capacity, attenuate inflammation, and promote lipid remodeling and utilization^6^. More recently, we and others have assessed chromatin accessibility in response to training^7,8^, demonstrating associations with transcriptional responses, as well as more rapid chromatin closure in trained compared to untrained murine skeletal muscle^9^. A few studies have also investigated post-translational modification (PTM) changes, including the phosphoproteome and acetylome^8,10,11^. Despite these efforts, limitations across studies include assessment of only one or two omics platforms applied in one sex or at a single timepoint. By extension, to our knowledge, no study has systematically and concurrently quantified the phosphorylation, acetylation, and cysteine oxidation modifications across the proteome, along with the transcriptome and epigenome in the same skeletal muscle sample, let alone examined male and female skeletal muscle responses in a temporal manner.

To address this gap, we leveraged and expanded upon the recently published MoTrPAC multi-omics data resource^8^ to characterize and compare the integrative molecular response in skeletal muscle to 1, 2, 4 or 8 weeks of progressive endurance exercise training in male and female rats. Specifically, our multi-omic approach integrated (with improved annotation using the updated rat reference genome) previously published metabolomic, epigenomic, transcriptomic and proteomic (global, phosphorylation) data, along with new lysine acetylation and cysteine oxidation (as a marker of redox state) data. Overall, our objective was to identify shared and sex-specific multi-omic molecular responses to endurance training in skeletal muscle, and relate them to phenotypic adaptations. These integrative multi-omic analyses aim to improve our understanding of the molecular drivers of the health benefits of exercise training in males and females, thereby offering translational potential to identify therapeutic targets for clinical development.

## Results

### Timewise multi-omic responses to endurance training in skeletal muscle

To comprehensively characterize the molecular response to endurance training, we subjected adult male and female rats to treadmill running for 1, 2, 4 or 8 weeks (1W, 2W, 4W or 8W), with sedentary (SED) animals as controls (Fig. 1A). A detailed description of the training protocol, along with phenotypic and cross-tissue multi-omic changes have been published^8,12^. In brief, endurance training induced expected physiological changes, as evidenced by an increase in VO_2_max after 8W (Fig. S1A), and increased *gastrocnemius* (GN) glycogen content and citrate synthase activity (Fig. 1B) ^12^. GN and *vastus lateralis* (VL), collected 48 hours after the last exercise bout, underwent global transcriptomics and metabolomics profiling, while GN was further analyzed for epigenomics, proteomics and multiple post-translational modifications (PTMs), including phosphorylation, acetylation and cysteine oxidation.

**Figure 1.**
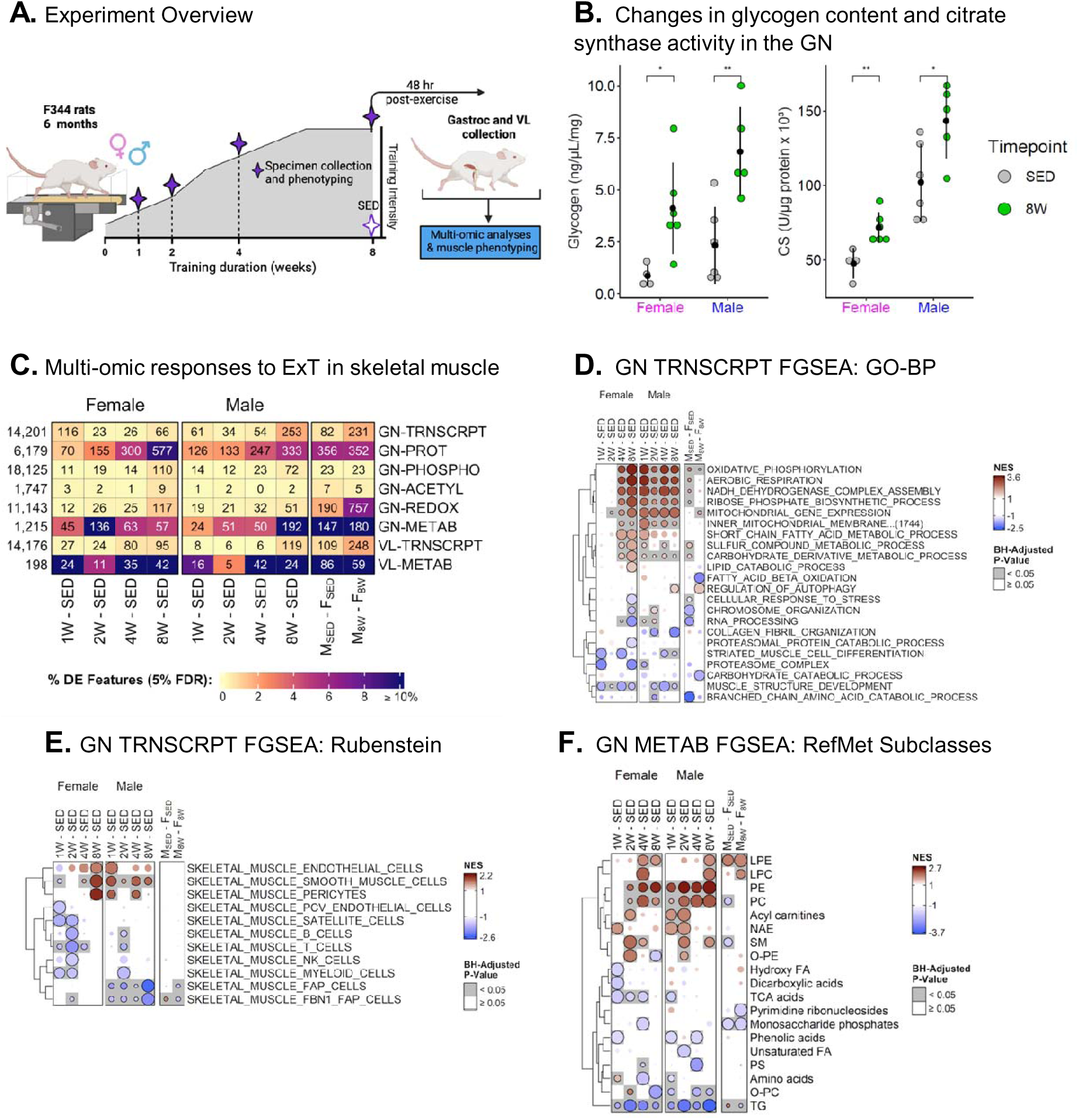
(A) Overview of 8W progressive endurance training protocol in male and female Fischer 344 rats. Muscles were collected from rats trained 1W, 2W, 4W, or 8W and SED controls (8W-aged matched) 48 hrs after the last exercise bout. This image was created with BioRender. (B) Representative dotplots of glycogen content (left) or citrate synthase activity (right) in gastrocnemius muscle (GN) of rats used for multiomics analyses. Data is grouped by sex and color coded by timepoint (SED or 8W). Brackets represent t-test p < 0.05 * or p < 0.01**. (C) Heatmap displaying differential features by sex and time point contrasts. The box color scale represents the percentage of significant DA features (BH-adjusted p-value <0.05) relative to the total number of detected features per ome (reported on the left), where each column represents a sex-specific contrast and each row represents a specific muscle and ome (reported on the right). (D) Heatmap representing timewise changes in Gene Ontology (GO) Biological Processes (BP) transcriptomic pathway enrichments by contrast in the GN muscle performed using FGSEA. (E) Heatmap of timewise changes of FGSEA gene set enrichments in the GN transcriptomics data curated using a reference gene set from skeletal muscle single cell transcriptomics data (Rubenstein et al.^13^). (F) Heatmap representing metabolomics pathway enrichments in the GN grouped according to RefMet subclass and analysed by FGSEA. In all bubble heatmaps (panels D-F), a black outline and gray shading indicates statistical significance (BH-adjusted p-value <0.05). In the sex-specific timewise contrasts, red indicates positive enrichment and blue negative enrichment, whereas in the M–F contrasts, blue indicates enriched in females and red indicates enriched in males. TRNSCRPT: transcriptomics; METAB: metabolomics; PHOSPHO: phosphoproteomics; PROT: proteomics; ACETYL: acetylomics; REDOX: cysteine oxidation proteomics.

We first assessed timewise changes across training groups and omes in the VL and GN (Fig. 1C). The VL largely displayed sex-conserved temporal increases in the number of differential transcripts and metabolites, with most changes occurring at later training timepoints. This was also observed for male GN, yet female GN displayed an earlier increase in the number of differential transcripts and metabolites: 1W and 2W, respectively (Fig. 1C; Fig. S1B). Across omes, the proteome (assessed only in the GN) displayed the greatest number of differential features at 8W in males and females (333 and 577, 5% and 9% of total features, respectively), with a progressive increase in the number of differential features over the endurance training period. After normalization for protein abundance, the PTM cysteine oxidation (redox) showed the greatest number of sex-biased differential features in GN after 8W, with more than 750 cysteine sites demonstrating sex differences in abundance at 8W (Fig. 1C; M_8W_ - F_8W_ contrast). Fewer sex-biased training responsive features were observed for protein phosphorylation and acetylation (Fig. 1C).

We next performed gene set enrichment analysis of the training-induced transcriptome (Fig. 1D-E and Fig. S1C-D). In GN and VL, males displayed an earlier (1W) enrichment in terms related to mitochondrial metabolism. In contrast, transcriptional enrichment of oxidative phosphorylation terms did not occur until 2W or 4W in GN and VL (Fig. 1D and Fig. S1C). Early enrichments in female GN and VL included decreased muscle structural development and proteasome pathways (Fig. 1D and Fig S1C).

Given the cellular heterogeneity of skeletal muscle, we next asked if multi-omic shifts arose from resident cell adaptation or altered cellular composition. Using a gene set enrichment approach with a human skeletal muscle single cell transcriptome dataset^13^, we estimated training-induced changes in cellular composition. Both sexes showed enrichment of transcripts indicative of endothelial and smooth muscle cells in the VL at 8W (Fig. 1E and Fig. S1D), and smooth muscle cells in the GN beginning in 4W females and 1W males (Fig. 1E), consistent with the pro-angiogenic stimulus of endurance training^14^. Additionally, males demonstrated a prolonged suppression of transcripts reflective of fibroadipogenic (FAP) cells in the GN and VL, which was prolonged relative to transient decreases observed in females (Fig. 1E). While endurance and resistance exercise promote FAP senescence and attenuate tissue FAP cell accumulation, sex differences in this response, to our knowledge, have not been previously reported^15,16^.

Metabolomics data revealed sex-conserved GN increases in plasmalogen species (PC and PE) and the PE-derived bioactive lipid species LPE, while triglyceride (TG) species decreased throughout training (Fig. 1F). Increases in PC and PE species have been reported with training, where a reduced PC:PE ratio has been linked to exercise-induced improvements in insulin sensitivity^17,18^. Fewer changes were observed in VL, likely due to the limited panel of metabolites analyzed in this tissue (Fig. S1E).

To fortify the translational relevance of this work, we compared our transcriptomic findings to a human 12-week intervention study^19^. This dataset was selected because of similar: 1) age of the cohort (i.e. young, adult), 2) sampling timepoint, 3) endurance training-induced increase in CS activity, and 4) both sexes were included. At the pathway level (using GO terms), the correlation between the male rat and male human enrichments was r = −0.16 (p-value < 2.2e^-16^). However, there was a relatively high correlation (Spearman’s r = 0.41; p-value < 2.2e-^16^) between the enriched pathways in female rats and female humans (Fig. S1G). Unexpectedly, the correlation of the response enrichments between female rats and male humans was much higher compared to that of male rats and male humans (Spearman’s r =0.42; p-value < 2.2e^-16^ compared to r=-0.16, Fig. S1F). Of note, the top 50 pathways with the highest average CAMERA-PR z-score across all comparisons were almost entirely composed of terms related to mitochondrial respiration (Fig. S1F, red points).

### Sex-specific chromatin accessibility changes affect transcriptional regulation of chromatin remodeling genes

To elucidate the regulatory mechanisms driving the transcriptional response to endurance training in skeletal muscle, we examined epigenetic modifications (ATAC-seq) and the relationships to differentially expressed genes. We identified 969 differentially accessible regions (DARs) across training timepoints (F-test adj. p-value < 0.1). Hierarchical clustering of the DAR magnitude of change (log2FC) revealed 5 distinct temporal patterns, each characterized by sex-specific response profiles (Fig. 2A top, and Fig. S2A).

**Figure 2.**
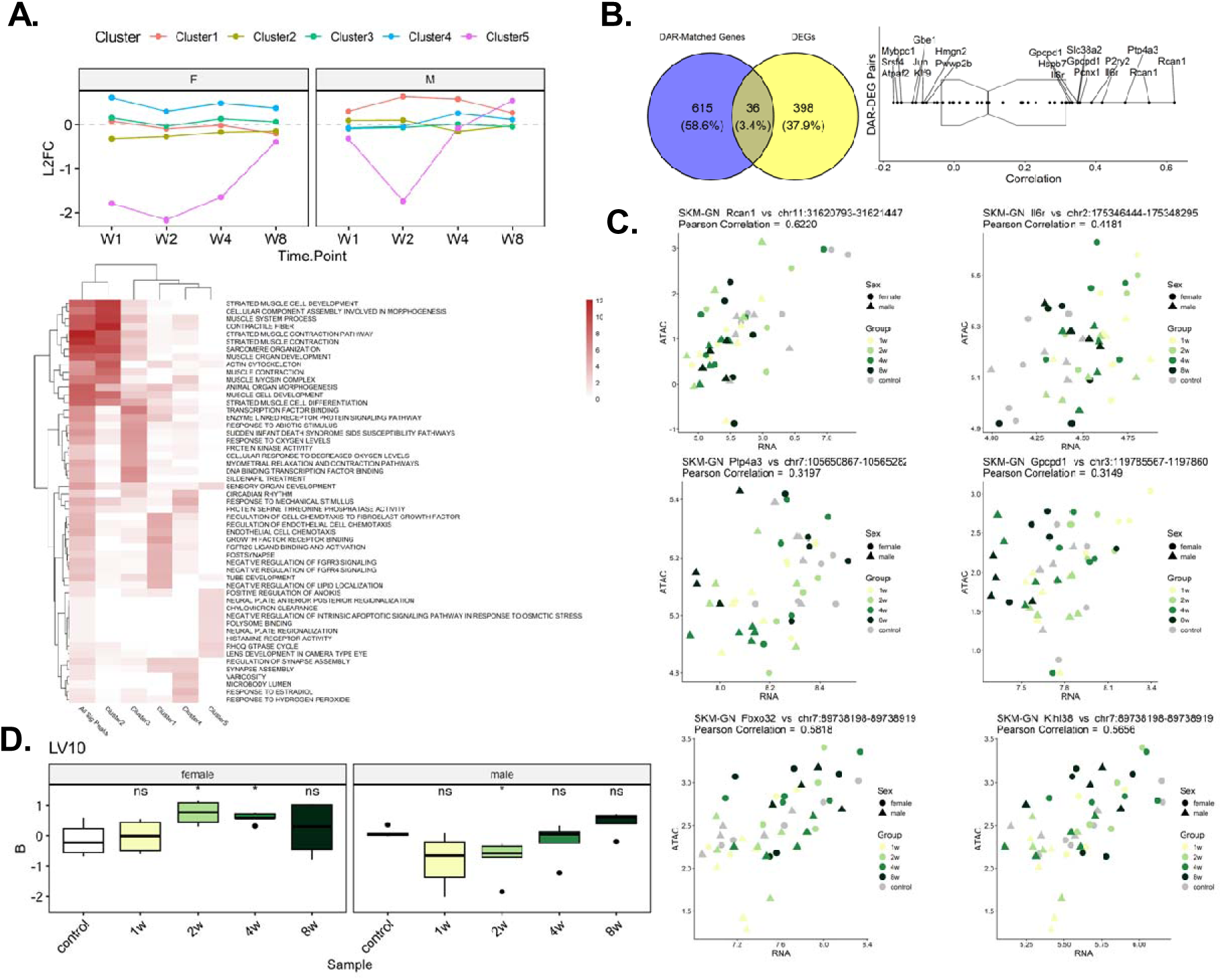
(A) Mean Log2FC trajectories for differentially accessible regions (DARs) aligned with each cluster identified in (Figure S2A) and their corresponding enrichments as performed using Over-representation analysis (ORA) for all DARs (all significant (Sig.) peaks) and for each DAR cluster as indicated below. (B) Venn diagram of overlapping DAR-annotated genes and DEGs (left) and representative correlation matrix with gene names (right). (C) Scatter plots of RNA and ATAC expression for DAR-DEG pairs including Rcan1, Il6r, Ptp4a3, Gpcpd1, Fbxo32, and Klhl38. Pearson correlation is labeled for each pair. Training timepoints and SED are labeled by color and sex is labeled by symbol. (D) Boxplot of LV10 pattern response to exercise training for each sex and time point. Significance labels represent within-sex changes relative to control: ns: p > 0.05, *: 5e-03 < p <= 0.05. B is a matrix value generated by PLIER to describe the expression pattern associated with that LV.

The five DAR clusters were associated with distinct biological processes (assessed using over-representation analysis, Fig. 2A, bottom). Clusters 1-3 were enriched for proximal promoter peaks (<= 1 kb from TSS) (Fig. S2B). Cluster 1, enriched for growth factor and muscle development genes and displayed male-specific training enrichments (Fig. 2A and S2A). Cluster 2, associated with muscle development and contraction terms and female-specific negative responses, while cluster 3 DARs, associated with transcription factor and oxygen sensing pathways, showed female-enriched training responses, although this cluster displayed the most variability in response. Synapse assembly and estradiol-associated cluster 4 DARs were upregulated in both sexes, although the male response was limited to later training timepoints.

Examination of overlap between DARs and differentially expressed genes (DEGs) across training timepoints revealed 36 DEGs (3.4%) with annotated DARs (Fig. 2B). Positively correlated DEG-DAR pairs included the calcineurin-inhibitor (*Rcan1*), the receptor for the myokine IL-6 (*Il6r*), protein tyrosine phosphatase 4a3 (*Ptp4a3*), and glycerophosphocholine phosphodiesterase 1 (*Gpcpd1*) (Fig. 2C). Applying Pathway Level Information ExtractoR, we identified latent variable 10 (LV10) reflecting highly correlated accessibility changes concentrated to a specific region of chromosome 7 (Fig. 2D). These peaks showed increased accessibility in female rats at 2W and 4W, and decreased accessibility in male rats at 2W, rebounding in both sexes at 8W, and overlapped or were located downstream of *Fbxo32* and its neighboring gene *Klhl38* (Fig. S2C-D), both of which showed highly correlated training responses with the LV10 peak set (Fig. 2C). *Fbxo32* and *Klhl38* govern muscle programs related to atrophy^20^, suggesting a potential effort by the muscle to coordinate transcriptional balance of muscle remodeling with exercise training.

### Endurance training drives alternative splicing of contractile genes

Beyond transcriptional changes, exercise training also drives alterations in RNA splicing that can impact gene isoform distribution and cellular processes^21^. Using a likelihood ratio test stratified by sex, we identified 19 isoforms corresponding to 13 genes that were affected by endurance training in the VL, whilst in the GN there were 13 differential isoforms corresponding to 7 genes (FDR < 0.05, Supp table X). Isoforms of *Eno3* (alternative 3’ splice site) and *Neb* (retained intron, Fig.S2E) were alternatively spliced in both tissues. GN-exclusive isoforms included *Tnnt3* (alternative 3’ splice site), *Nr4a1* (alternative first exon) and *Myh1* (alternative 5’ splice site, A5SS), while VL-exclusive isoforms included *Fth1* (A5SS), *Myl1* (retained intron and A5SS), and *Sdhc* (retained intron) (Fig.S2E, Supp table X). Isoforms were enriched for structural constituents of muscle, muscle contraction, muscle development, alpha actinin binding, and contractile fibers in both GN and VL (Fig. S2F). These spliced sarcomeric gene isoforms have been suggested to play differential roles in myogenesis and contraction, with potential implications for adaptations to endurance exercise^22^.

### Multi-omic timewise response trajectories show earlier induction of mitochondrial pathways in females

Given the sex-specific temporal patterns observed for individual omes, we next applied an integrative analysis to characterize multi-omic response trajectories. Henceforth our analyses focus on GN, which underwent the most extensive multi-omic profiling. To systematically identify timewise response trajectories, we applied C-means clustering across the transcriptome, proteome and metabolome, identifying 12 response trajectories (Fig. S4A-B), to reduce the dimensionality of findings into biologically interpretable context, we then focused on four clusters with distinct temporal trajectories observed in both sexes (Fig. 3A). Non-parametric pre-ranked Correlation Adjusted MEan RAnk (CAMERA-PR) was performed to assign biological relevance to these clusters.

**Figure 3.**
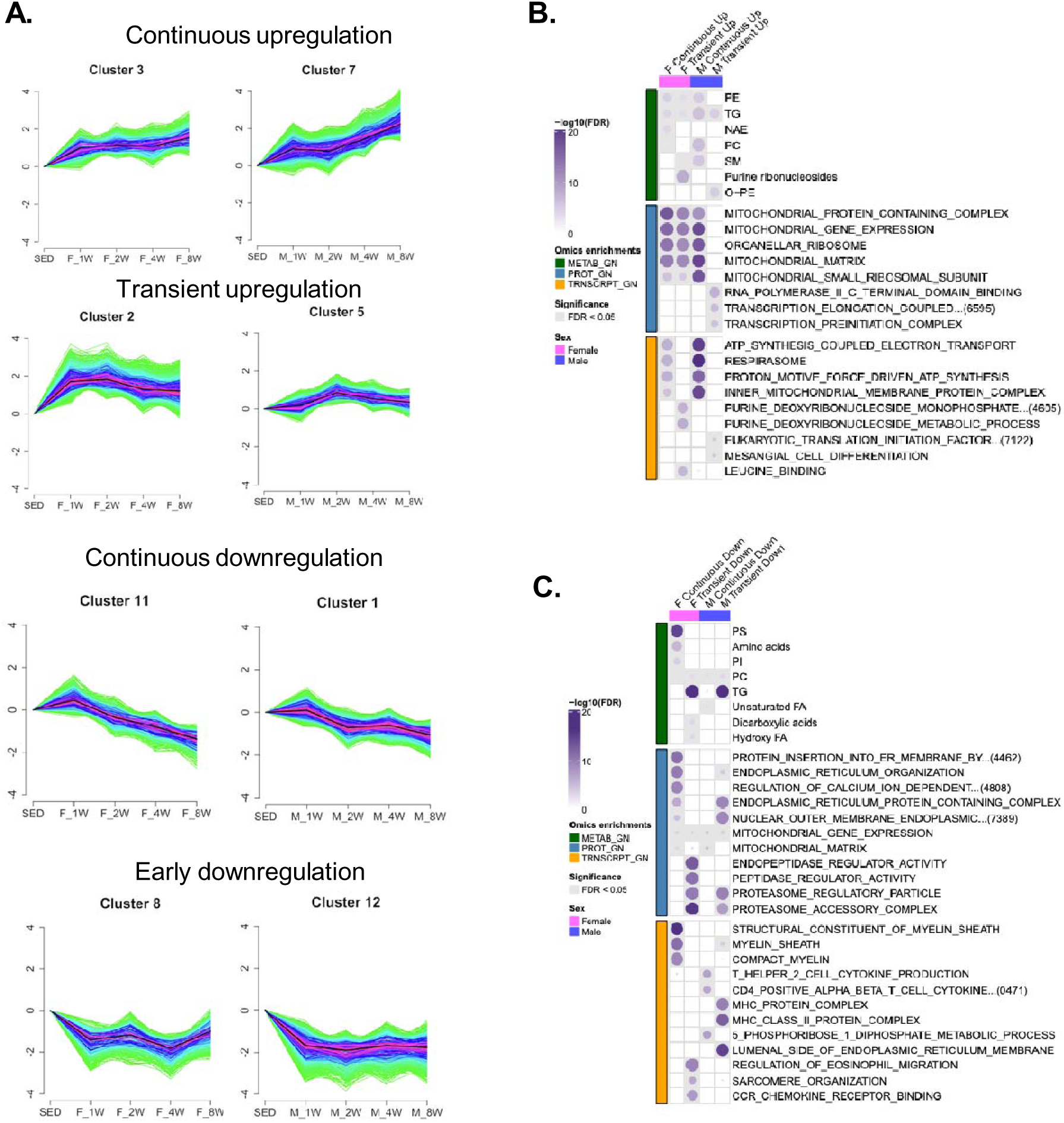
(A) Select multi-omic Fuzzy C-means trajectories for GN metabolomics (METB), transcriptomics (TRNSCRPT), and proteomics (PROT) molecular features in females (left panels) and males (right panels), where the dark shading indicates the distribution centroid. A qualitative description of the trajectory male/female pair is provided above for context. (B) ORA enrichment analysis for features localizing to female and male clusters displaying continuous or transient upregulation. Leftward colors represent enrichments per ome, where METAB: RefMet enrichments, and TRNSCRPT and PROT: GO enrichments. Gray shading indicates adjusted p-value <0.05. (C) ORA enrichment analysis for features localizing to female and male clusters displaying continuous or early downregulation. Leftward colors represent enrichments per ome, where METAB: RefMet enrichments, and TRNSCRPT and PROT: GO enrichments. Gray shading indicates adjusted p-value <0.05.

Of the clusters displaying progressive increases over time (female cluster 3, a male cluster 7) were enriched for mitochondria-associated transcripts and proteins as well as PE, PC and TG metabolites (Fig. 3A-B; Fig. S3C-D). Among clusters displaying transient upregulation between 1W-4W (female cluster 2, male cluster 5), the female cluster displayed metabolomic enrichments in purine ribonucleosides, transcripts related to leucine and amino acid binding and deoxyribonucleosides, and mitochondrial proteins (Fig. 3B and Fig. S3C-D). The corresponding cluster in males displayed transcriptomic and proteomic enrichments in transcription and translational terms (Fig. 3B and Fig. S3C-D). Of the clusters displaying continuous downregulation (female cluster 11, male cluster 1), both displayed metabolomic enrichment in PC species, with other notable sex differences. In females, cluster 11 displayed enrichments in myelin sheath transcriptomic terms and proteomic features related to mitochondrial gene expression, whereas male cluster 1 was predominantly enriched in immune-related transcripts (Fig. 3C and Fig. S3C-D). In both sexes clusters displaying early and suppressed downregulation (female cluster 8, male cluster 12), were enriched for proteasomal proteins (Fig. 3C and Fig. S3D). Together our findings identify broadly sex-distinct temporal dynamics of the progressive response to endurance exercise training, with early downregulation of proteasomal proteins being a sex-conserved training response.

### Sex-divergent post-translational modifications reveal opposing redox states in trained muscle

PTMs are an important means of regulating protein function, which motivated our examination of temporal, sex-dependent changes in protein PTMs, corrected for protein abundance. Overall, PTMs showed a delayed response to endurance training with the majority of changes occurring at 8W, despite temporal increases in global proteomic features (Fig. S4A-B and D). Largely, global proteomics revealed sex-conserved increases in mitochondrial terms, consistent with c-means analyses (Fig. 3A-B), albeit females displayed an earlier (1W and 2W) increase in mitochondrial terms in females compared to males (4W and 8W, Fig. 4A). At 8W, males displayed enrichment in terms related to response to heat and humoral immunity, whereas females displayed enrichments in lipid oxidation processes. Examination of phosphoproteomic responses displayed little overlap between sexes, with 13 phosphosites responding to 8W training in both sexes (Fig. S4B). Among the most sexually dimorphic phosphoproteins at 8W, PRKN S77 phosphorylation was higher in females; this site is adjacent to its PINK-mediated ubiquitin-like domain that promotes mitophagy^24^ (Fig. S4C), suggestive of female-specific mechanisms of muscle remodeling. Males displayed elevated stress-associated MAPK14 (MAPK p38L) S28 phosphorylation; while the function of this site is unknown, phosphorylation of an adjacent site, S38, promotes myogenesis^25^.

**Figure 4.**
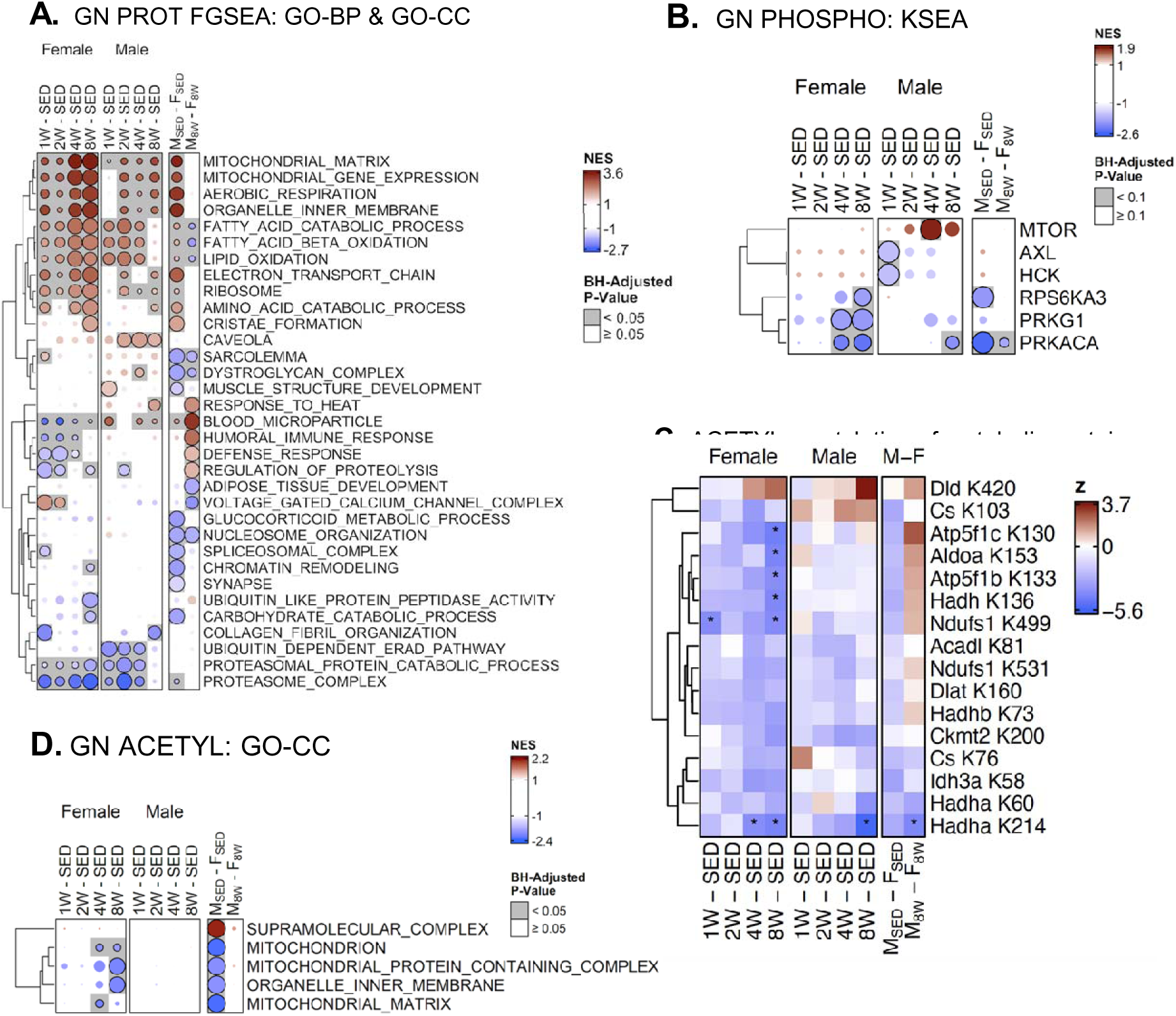
(A) Heatmap displaying GN GO-BP and GO-Cellular Component (CC) proteomic enrichments by contrast using FGSEA gene set enrichment analysis; a black outline and gray shading indicates statistical significance for BH-adjusted p-values <0.05. (B) Heatmap displaying GN kinase-substrate enrichment analysis (KSEA) results from phosphoproteomics data. A BH-adjusted p-value threshold of p<0.1 was used for analyses, indicated by a black outline and gray shading. (C) Feature-level heatmap depicting the top differentially regulated acetylation modifications across metabolic proteins from the timewise contrasts;* indicates significance at BH-adjusted p-value <0.05 per feature in DA model. (D) Heatmap displaying the results of FGSEA analysis of the GN acetylproteomic dataset, against the GO-CC database.

Despite limited overlap in individual phosphosites, several sex-conserved phosphorylation changes were linked to muscle remodeling processes. These included the mTOR inhibitor DEPTOR (S244), the heat shock protein family member HSPB1 (Y137) and phosphorylase kinase regulatory subunit 1A (PHKA1 S1019) (Fig. S4C). Notably, PHKA1 showed C-terminal serine dephosphorylation in both sexes. Since PHKA1 links calcium signaling with glycogen breakdown during muscle contraction, and PHKA1 S1018 is linked to glycogenolysis^26,27^, our finding suggests reduced reliance on muscle glycogen during exercise following training. A regulatory subunit of myosin phosphatase (PPP1R12B, or MYPT2) displayed sex-conserved dephosphorylation of S502; an adjacent site (S504) is phosphorylated following insulin exposure^28^. Further, CSRP3 S95 phosphorylation increased with training; CSRP3 (Muscle LIM protein) facilitates autophagosome formation in myogenesis, yet its phosphorylation has not been documented in the skeletal muscle with exercise and suggests a mechanism promoting muscle protein quality control with training^29^.

To gain further insight into the potential drivers of phosphoproteomic changes with training, we performed kinase-substrate enrichment analysis using PhosphoSitePlus^30^. Two kinases displayed sex-conserved decreases at 8 weeks; RPS6KA3, which is associated with stress responses, and the cAMP-dependent protein kinase catalytic subunit alpha, PRKACA (Fig. 4B). In contrast, mTOR was enriched only in 4W males, and attenuated by 8W. Notably, PRKACA displayed the largest sex bias, being higher in SED and 8W females (Fig. 4B).

Despite few protein acetylation modifications, both sexes displayed deacetylation of HADHA K214 (Fig. 4C), a key protein for fatty acid oxidation, which we previously reported to be affected by training in other metabolic tissues, including the heart and liver^31^. This finding is consistent with the induction of Sirtuin 3 (SIRT3) protein, a mitochondrial deacetylase that targets HADHA^32,33^, in both males and females (Supplemental Table X). Gene set enrichment analysis revealed coordinated, pathway-level deacetylation of mitochondria-associated proteins in 8W females but not males (Fig. 4D), driven by deacetylation of complex I (NDUFS1 K499) and V proteins (ATP5F1C K130 and ATP5F1B K133) (Fig. 4C).

Cysteine oxidation proteomic analysis revealed marked sex differences in the training response (Fig. 5A-C; Fig. S5A). In 8W males, multiple mitochondrial pathways including oxidative phosphorylation, inner mitochondrial membrane, TCA cycle and cristae formation showed increased cysteine oxidation (Fig. 5C and Fig. S5A). In contrast, females increased oxidation of proteins associated with carbohydrate metabolism and reduced oxidation of proteins involved in lipid and BCAA metabolism (Fig. 5C-F; Fig. S5A-B), resulting in sex-specific differences at 8W. These 8W sex-specific differences in proteins related to cristae formation in females relative to males (Fig. 5E), may impact electron flow in the ETC and available surface area for cellular respiration^34^.

**Figure 5.**
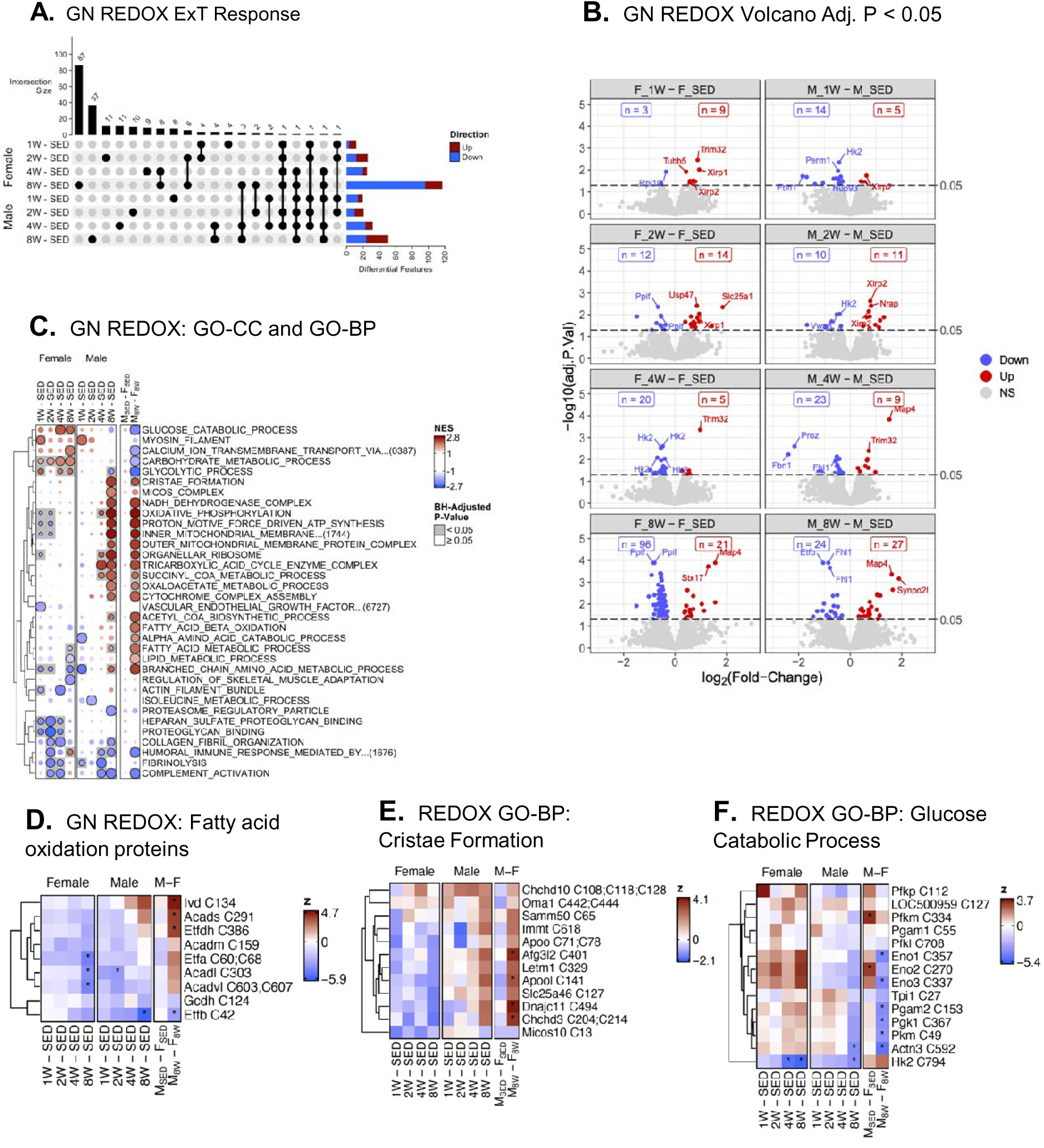
(A) UpSet plot displaying the number of significantly different cysteine oxidation protein modifications per training timepoint in female and male rats. A line indicates shared DA features per contrast in males and/or females, where the number of shared features is indicated as intersection size (top) and the number of DA features per timepoint and sex is indicated on the right. (B) Volcano plots showing the results of statistical analysis of cysteine oxidation proteomics at each training timepoint in males and females. Statistically significant sites (adjusted p-value <0.05) are labeled in red (increased cysteine oxidation) or blue (decreased cysteine oxidation) and the number of significant sites in either direction are reported within each panel. The top 5 most significant features within each panel are labeled with their corresponding gene IDs. (C) Heatmap displaying the results of FGSEA analysis of the GN cysteine oxidation proteomics dataset, against the GO-BP and GO-CC databases; terms were manually curated to reduce redundancy and show pathway enrichments pertaining to different biological functions. (D-F) Feature-level heatmaps showing cysteine oxidation of select proteins involved in fatty acid oxidation (D), cristae formation (E), and glucose catabolic processes (F); * indicates significance at BH-adjusted p-value <0.05.

Several sex-conserved changes were also observed. Four-and-a-half LIM protein 1 (FHL1), a protein implicated in skeletal muscle hypertrophy and NFATc signaling^35^, showed decreased oxidation at multiple sites (Suppl table X). Hexokinase 2 (HK2) and myosin heavy chain 2 (MYH2) also displayed sex-consistent reduced oxidation at 8W (Fig. S5C). Oxidation of MYH2 reduces contractile force in skeletal muscle of rats with chronic heart failure ^36^ and in response to muscle unloading ^37^, alluding to a potential regulatory role of training-induced shifts towards a more reduced state of sarcomeric proteins, a mechanism that, to our knowledge, has not been explored previously.

Integration of global and PTM proteomics further elucidated sex differences in mitochondrial protein regulation with training. Females more commonly upregulated Complex II, III, and IV proteins, whereas Complex I and V proteins displayed largely sex-conserved responses (Fig. 6A; S6A-E)^38^. These adaptations are congruent with our findings of diminished mitochondrial cysteine oxidation. Supporting a greater proclivity for female skeletal muscle to form mitochondrial supercomplexes, a key stabilizer of Complex III and IV stability, supercomplex assembly factor COX7A2L, displayed female-specific increases at 8W^39^ (Fig. S6D). Training also induced enrichment of several proteins related to cristae structure and mitochondrial fusion and fission in female skeletal muscle (Fig. 6B), indicative of more efficient energy turnover in response to training. To support this, females generally displayed GN nucleotide patterns indicative of energetic efficiency relative to males at 8W (Fig. 6C). However, both sexes displayed training reductions in the PC:PE ratio (Fig. 6D), albeit 8W females did not reach significance driven by an outlier, indicative of mitochondrial membrane remodeling to favor energetic efficiency^18^. Overall, these findings suggest that while mitochondrial pathways are enriched in both sexes, female-biased increases in cristae architectural proteins with training may reduce mitochondrial oxidative stress.

**Figure 6.**
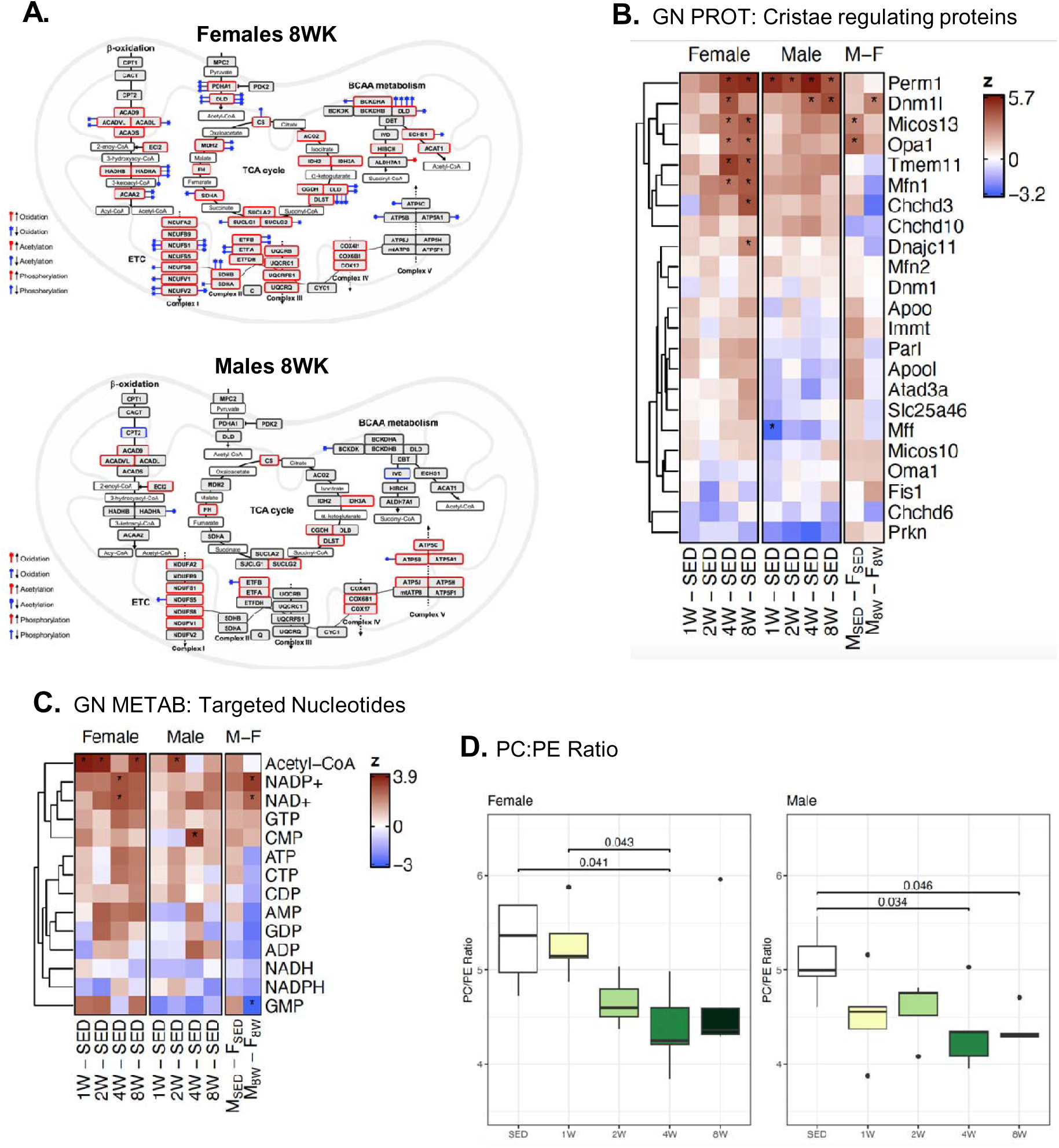
(A) Schematic displaying 8W changes in female (top) and male (bottom) mitochondrial proteins and PTMs. A red square indicates a significant increase in global protein abundance in 8W male or female relative to SED, whereas a blue square indicates reduced protein abundance. PTMs to mitochondrial proteins are indicated by defined symbols (left key); color indicates significant change in PTM on named protein relative to normalized protein abundance (blue=decrease, red=increased). (B) Feature-level heatmaps displaying the z-score of GN cristae-related protein changes per contrast; * indicates significance at BH-adjusted p-value <0.05 per contrast. (C) Effects of training and sex on targeted nucleotide metabolite abundance in GN; * indicates significance at BH-adjusted p-value <0.05 per contrast. (D) Boxplots displaying timewise changes per sex in PC:PE lipid ratio using internal standards. Tukey HSD test was performed as a post hoc analysis following ANOVA to identify which pairwise comparisons were statistically significant; statistical significance (adjusted p-value) is indicated above the contrast pairs, where each training timepoint is compared to SED.

### Proteomic modules link mitochondrial remodeling to aerobic capacity gains

To identify multi-omic features most closely associated with physiological adaptations, we performed Weighted Gene Co-expression Network Analysis utilizing global proteomics and then correlated resultant Module eigengenes (MEs) to 14 different phenotypes (Fig. 7A-B). Including sex in the model yielded the highest number of phenotype–ME correlations, suggesting sex-biased regulation of the proteome. Male sex was highly correlated to ME-turquoise (mitochondrial), which was likely driven by baseline enrichments (Fig. 4A) as features localized to this ME increased in both sexes with training, and it was positively correlated to increases in VO_2_max, lean mass, and muscle citrate synthase activity (Fig. 7A and C; Fig. S7A-B). ME-red (glycogen metabolism and calmodulin binding) displayed enrichments in females, yet tended to decrease in both sexes with training, which is likely reflective of reduced dependence of glycogen as it was negatively correlated to glycogen and CS. ME-green displayed the most sex-biased trajectories, increasing in males and decreasing in females; it was enriched in actin binding and hemostasis terms (Fig. 7A and C).

**Figure 7.**
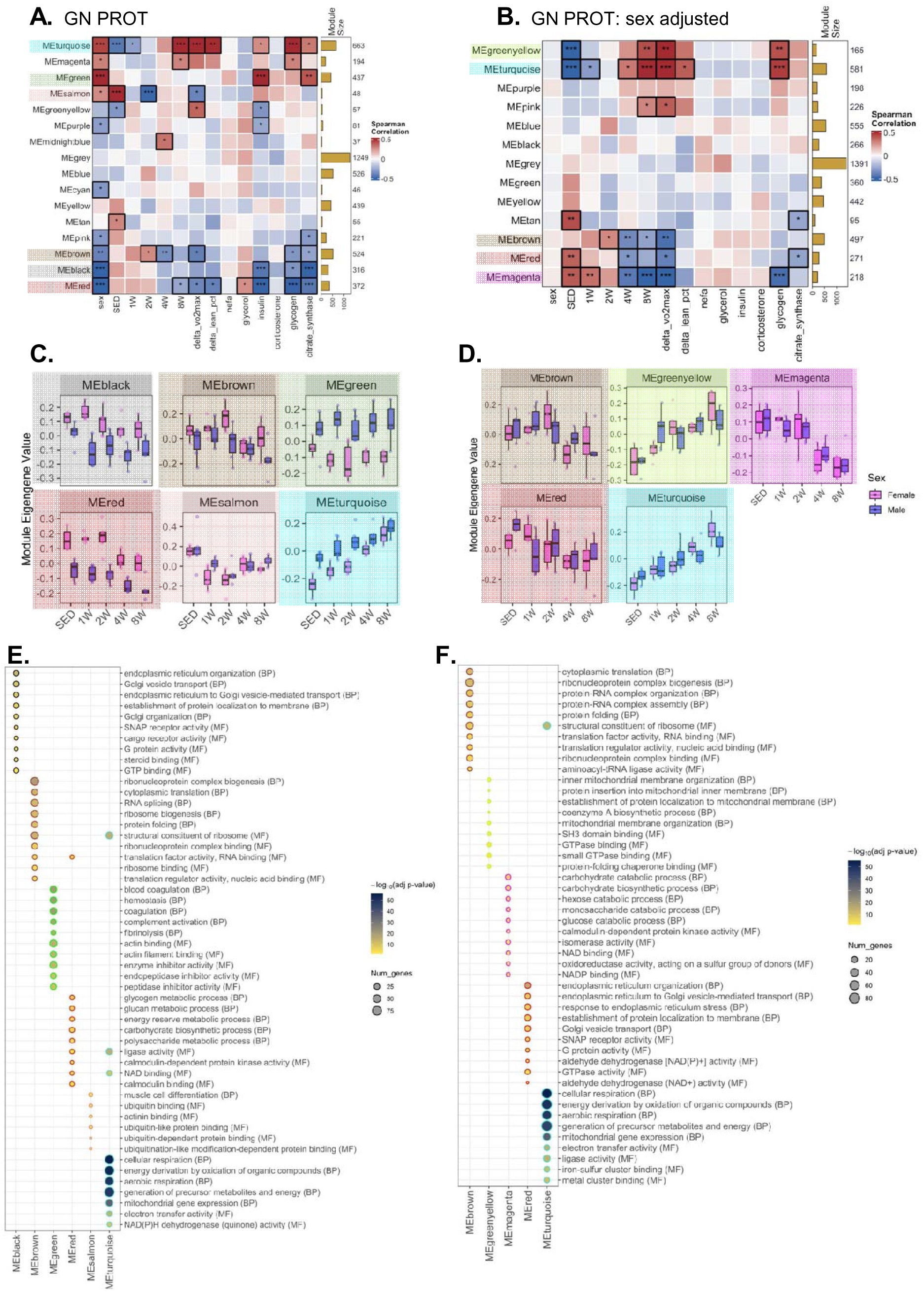
(A-B) Heatmaps showing the Spearman correlation between module eigengenes (MEs) generated through weighted gene co-expression network analysis (WGCNA) and various clinical and physiological measures for proteomics including sex as a phenotypic variable (A) and after adjustment for sex (B). (C-D) Over-representation analysis (ORA) of the proteins contained in select WGCNA MEs generated without (C) and with (B) adjustment for sex.

Given sex was the predominant correlate to proteomic MEs, we re-ran the model adjusting for sex to gain insight into sex-conserved proteome-phenotype relationships (Fig. 7B and S7C-D). Two modules, ME-greenyellow and ME-turquoise, both with mitochondrial enrichments, were positively correlated to increases in VO_2_max and glycogen (Fig. 7B and D). Conversely, the brown, red, and magenta MEs decreased at 8W, were negatively correlated to VO_2_max and enriched for features related to cytoplasmic translation, endoplasmic reticulum, and carbohydrate catabolic processes, respectively (Fig. 7C and E), respectively. Together, these findings suggest a partitioning of proteomic pathways to favor mitochondrial function and fatty acid oxidation linked to training improvements in both sexes.

## Discussion

By integrating newly generated and publicly available multi-omic data from male and female rats, we provide the most comprehensive multi-omic map of skeletal muscle remodeling in response to endurance training to date. We find that while males and females share critical adaptations, including increased mitochondrial capacity and reduced proteasome activation, significant sex differences are evident in the temporal dynamics of adaptations and in post-translational modifications of metabolic proteins. Cysteine oxidation proteomics data uncovered previously unrecognized sex-divergent cysteine oxidation changes with training, with females showing responses indicative of improved mitochondrial redox buffering capacity, whereas males exhibit attenuated protein turnover signaling. Together, our findings establish a foundation for understanding how biological sex influences skeletal muscle adaptations to endurance exercise training.

The temporal dynamics of the transcriptomic and proteomic responses to training differed between males and females. Following training, both sexes displayed enrichments in mitochondrial genes and proteins, with females displaying an earlier (1W and 2W) increase in mitochondrial protein enrichments. Trajectory analysis identified a transient enrichment in ribosome-associated transcripts in females that may have facilitated accelerated protein translation of mitochondrial genes despite delayed transcriptional enrichments. Estrogen-mediated effects on translational stability^40^, or sex differences in the rate of mitochondrial protein turnover^41^ could also contribute. The transcriptomics data was also used for cell type deconvolution, identifying sex-consistent changes in skeletal muscle cellular composition with training, including enrichment of endothelial and smooth muscle cells, consistent with increased angiogenesis with training.

Integrative cysteine oxidation and global proteomic analyses revealed pronounced sex-specific metabolic protein PTMs that varied by subcellular localization. Consistent with human studies^42^, males exhibited increased oxidation of mitochondrial proteins with training, whereas females showed decreased oxidation across several mitochondrial proteins—most notably those of complex I and V, key components of mitochondrial supercomplexes. These cysteine oxidation patterns coincided with female-specific increases in the abundance of complex I–IV proteins and factors governing cristae ultrastructure, stability, and fusion, including CHCHD3, COX7A2L, OPA1, and MFN1^31,43^. Together, these findings support a model in which enhanced supercomplex formation improves mitochondrial efficiency by buffering electron leak and reducing reactive oxygen species formation at the ETC.

Intriguingly, despite decreased cysteine oxidation of mitochondrial and fatty acid–related proteins, females displayed greater oxidation of glycolytic proteins following training, generating sex differences evident by 8 weeks. This observation suggests sex-biased localization of reactive oxygen species. Overall, the delayed PTM response, predominantly occurring after 8 weeks of training in both sexes, suggests that post-translational modifications represent a mechanism to further optimize functional improvements beyond initial increases in global protein abundance. Both sexes also displayed decreases in the ratio of the phospholipid species PC:PE with training. Importantly, the composition of PE and PC in skeletal muscle is linked to improved metabolic health, whereby both increase with endurance training, also in human skeletal muscle, with PE increasing to a greater extent, thus decreasing the PC:PE ratio^18^.

Males reduced molecular profiles linked to protein turnover with training. Cell culture studies have demonstrated testosterone enhances mTOR activity and promotes skeletal muscle protein synthesis^44^ and, we observed elevated mTOR signaling specifically in males after 4 weeks of training. By eight weeks, males also showed decreased phosphorylation of OPTN S342 and PRKN S77 suggesting reduced Parkin-dependent mitophagy ^49,50^. In contrast, estrogen signaling supports mitochondrial biogenesis, oxidative metabolism, and activation of autophagyLrelated pathways, providing a mechanistic basis for the earlier engagement of targeted protein qualityLcontrol processes in females ^45,46^. In line with this, females displayed increases in PRKN S77 phosphorylation. As a key inducer of PINK1-mediated mitophagy, this indicates enhanced mitochondrial clearance in females in response to endurance training.

Several lines of evidence support enhanced and earlier protein quality control in females. Females exhibited BAG3 S294 (human S291) phosphorylation associated with increased CAA^47^, along with early and transient chromatin accessibility changes for *Fbxo32* and *Klhl38*, two genes involved in ubiquitin-mediated protein turnover^20,48^. These chromatin changes were paired with increased gene expression, indicating coordinated epigenetic and transcriptional activation of ubiquitin-mediated proteolysis. Interestingly, males show reduced chromatin accessibility at the *Fbxo32/Klhl38* loci at two weeks, an example of sex-divergent regulation of the epigenome. Overall, these observations suggest that males favor anabolic signaling and delay selective protein turnover, whereas females appear to prime these pathways earlier in training, perhaps as a quality control mechanism to promote mitochondrial biogenesis and efficiency ^51^.

Several limitations warrant consideration. The cohort size (N=6 per sex and timepoint), while sufficient for multi-omic profiling of rat training effects, may limit detection of more subtle effects. The sampling timepoint of 48 hours after the last exercise bout was selected to minimize acute effects, yet some transient responses may remain. Although the human cohort utilized in this study was selected to be reasonably comparable to the rats with regards to age, sex, sampling timepoint and physiological response, further integration of human and murine data sets are needed to identify ideal models for skeletal muscle studies. Future studies examining the impact of age and sex hormones will facilitate further understanding of sex-biased adaptations and their implications for muscle function across the lifespan and in disease states.

Overall, our findings demonstrate that biological sex profoundly affects the molecular response to endurance exercise training at multiple regulatory levels. Females display proteomic and post-translations adaptations indicative of improved mitochondrial efficiency, characterized by enhanced supercomplex formation, reduced mitochondrial cysteine oxidation, and early activation of protein quality-control mechanisms. Males, despite earlier induction of mitochondrial transcripts, show a more modest increase in mitochondrial protein abundance, but a greater oxidation of mitochondrial proteins in response to training. This data supports investigation of molecular features across sexes and regulatory levels, towards better understanding skeletal muscle adaptations to exercise training, and establishes a foundation for more precise training interventions and identification of therapeutic targets.

## Methods

### Experimental Design

The experimental design has been previously described in detail ^52^. In brief, six-month-old male and female Fischer 344 rats performed progressive treadmill exercise training five days per week for 1, 2, 4, and 8 weeks, or remained sedentary. Tissues, including *gastrocnemius* and *vastus lateralis* muscles were collected 48 hours after the last exercise bout. The physiological effects of the endurance exercise training were assessed by VO_2_max testing for aerobic capacity, NMR for body composition, as well as muscle specific measures of glycogen content and citrate synthase (CS) enzyme activity. Phenotypic adaptations have been described in detail elsewhere^12^. All animal procedures were approved by the Institutional Animal Care and Use Committee at the University of Iowa.

### Multi-Omic Data Generation and Processing

Full details of the methods used for sample processing, data collection, data processing, data normalization and batch correction for transcriptomics, proteomics, phosphoproteomics, and metabolomics/lipidomics have previously been described in detail elsewhere ^8^. These methods, as well as methods for cysteine oxidation proteomics sample processing and data processing, are described in the Supplementary Methods file.

### Differential Analysis

RNA-Seq data was analyzed using the *edgeR*^53^ and *limma* ^54^ R/Bioconductor packages, following the workflow described by Law *et al*. ^55^. Briefly, low-abundance transcripts were removed from the raw count data with *edgeR::filterByExpr* to more accurately estimate the mean–variance relationship in the data. Then, a multidimensional scaling (MDS) plot was generated from the log_2_ TMM-normalized counts per million reads to visualize samples. Two samples from the vastus lateralis and three from gastrocnemius were identified as outliers in the principal component analysis and MDS plots and removed ^8^ (sample IDs are reported in the “Data Processing” section of the Supplemental Methods). After outliers were removed, differential analysis of -omics data was performed as described previously ^56^, and detailed below, with the following minor modifications: 1) *edgeR::voomLmFit* was used instead of *limma::voomWithQualityWeights* for the analysis of transcriptomics data, and 2) the mean–variance trend was not estimated separately for each metabolomics platform. Additionally, the comparison of males to females at 8W was included in addition to the SED comparison.

Analysis of cysteine oxidation proteomics and acetyl proteomics data was performed the same as for the phosphoproteomics data, while analysis of ATAC-Seq data was performed the same as for the transcriptomics data.

Additionally, RNA integrity number, median 5’-3’ bias, percent of reads mapping to globin, and percent of PCR duplicates as quantified with unique molecular identifiers were included as covariates in the RNA-Seq model after they had been mean-imputed and standardized ^52^. For each -ome, we specified a no-intercept model containing the experimental group (each combination of sex and timepoint; factor with 10 levels) and any covariates. Then, robust Empirical Bayes moderation was carried out with *limma::eBayes* to squeeze the residual variances toward a common value (RNA-Seq only) or a global trend (proteomic datasets)^57^. To control the false discovery rate (FDR), p-values were adjusted across sets of related comparisons by -ome using the Benjamini–Hochberg (BH) procedure.

### Differential Splicing Analysis

Isoform abundance was quantified from the RNA-Seq data with the *SGSeq* ^58^ R package, which quantifies alternative splicing events—identifying transcript start, end, or internal gene splicing events. Isoform quantification was performed using the *SGSeq::analyzeFeatures* function, with filtering criteria requiring an average percent-spliced-in (PSI) of at least 10% across samples.

Transcript isoforms were filtered to include only those with an average of at least five read counts across all samples. For each gene, the proportion of reads corresponding to each transcript isoform was calculated as the percentage of total counts for that gene within each sample. Percentages were then logit-transformed to stabilize variance and avoid boundary effects from proportions. In cases where a gene contained only two isoforms, only the first isoform was analyzed, as the second isoform’s percent-spliced-in value is statistically redundant (i.e., 100% – the first isoform’s proportion). Extreme logit values were capped at 5 or −5 to avoid infinite transformations.

To evaluate if a transcript isoform is alternatively incorporated due to exercise training, a likelihood ratio test (LRT) was performed for per isoform to test whether incorporating sex and exercise timepoint information improved model fit. For each isoform’s logit-transformed percent-spliced-in values, a full linear model including group and timepoint as predictors was compared against a nested model containing only the intercept. Pathway characterization of alternatively spliced genes was conducted via CAMERA-PR gene set enrichment analysis as described below, with each gene represented by the isoform with the highest F-statistic.

### Gene Set Testing

We performed three complimentary approaches to gene set enrichment. Fast gene set enrichment analysis (FGSEA) ^59^ was implemented using the *fsgea::fgsea*^59^ function. Pre-ranked Correlation-Adjusted MEan-RAnk (CAMERA-PR) gene set testing was performed with the *limma::cameraPR* ^54^ function in R to test for consistent up- or down-regulation or post-translational modification of gene sets ^60^. For both approaches, the Z-Score-transformed moderated t-statistics from the differential analysis results were used as input. We additionally performed overrepresentation analysis using Fisher’s exact test, as implemented in the clusterProfiler ^61^. package, to characterize specific subsets of features. The significance thresholds and selection criteria used for these analyses are detailed per analysis.

For transcriptomics (including differential splicing) and proteomics (with exception of phosphoproteomics), the gene sets tested with consisted of Gene Ontology Biological Process terms from the human Molecular Signatures Database (MSigDB <u>v2023.1.Hs</u>) ^62–64^. For phosphoproteomics, kinase curations were made using curations from PhosphoSitePlus ^65^. These gene sets were processed further (including converting human genes to their rat orthologs), and this is described in the Supplementary Methods (*Processing of Biomolecular Sets*).

### Multi-Omics Analysis PLIER

To delve into temporal trends in the chromatin accessibility response to exercise training, we applied PLIER ^66^, which takes an expression matrix of features vs samples as input and outputs patterns of activity, in the form of latent variables (LVs), and connects these patterns with user-provided prior information, such as reactome or KEGG pathways. ATACseq peaks were annotated to their closest gene TSS and gene-specific pathway enrichments as well as known binding sites for transcription factors within each peak from the HOMER rat TF database ^67^. We use the PLIER package R function *num.pc* to identify the number of significant principal components for the singular value decomposition (SVD) of the data and specify double this number as the desired LV count. PLIER’s prior knowledge input was set to be the C2 collection of MSigDB. *num.pc* calculated 14 significant principle components for the SKM-GN ATACseq data resulting in PLIER generating 28 LVs that best account for all patterns identified in the dataset, optimizing individual LV trajectories over the samples and feature associations to minimize the difference between normalized input data and PLIER LV output, while also accounting for feature associations with prior knowledge in the form of pathway enrichments. Significance of an LV to differentiate exercise training from control at a given time point is determined using the *stat_compare_means* R function from the ggpubr package.

### WGCNA

Weighted Gene Co-expression Network Analysis (`WGCNA`; R package v1.73; Langfelder & Horvath, 2008) was used to identify modules of co-expressing proteins associated with exercise training and key physiological and biochemical traits. Protein expression data (6,179 proteins) were filtered using the `goodSampleGenes()` function to exclude 392 proteins with excessive missing or zero-variance values; all samples passed quality control. Proteins with >30% missing values were further removed, and remaining missing values were imputed using k-nearest neighbors (KNN) method, resulting in a final data matrix of 57 samples and 5,265 proteins. For sex-adjusted analyses, protein expression values were regressed on sex and residuals were used for network construction. Co-expression networks were built using `blockwiseModules()` with a signed network (`networkType = “signed”`), a soft-thresholding power selected by `pickSoftThreshold()` to optimize scale-free topology (R² > 0.85), and minimum module size of 20 proteins. Module eigengenes (first principal component of each module) were computed for downstream analysis. Proteins assigned to the “grey” module (MEgrey) represent those whose expression profiles did not correlate strongly with any specific module and therefore were not included in co-expression clusters. Module–trait associations were evaluated via Spearman correlation; p-values adjusted for multiple testing using the Benjamini-Hochberg method. Functional enrichment was performed for each module using `enrichGO()` (`clusterProfiler` v4.12.6; Wu et al., 2021) against the rat genome annotation (`org.Rn.eg.db` v3.19.1). Significant GO terms were defined as those with Benjamini-Hochberg adjusted p-value < 0.05.

### Comparison to Human Data Sets

Human RNA-seq count data from the EpiTrain (“Epigenetics in Training”) study were used to identify exercise-responsive genes ^19^. Briefly, male and female young sedentary volunteers underwent supervised one-legged knee-extension exercise training for 3 months (45 minutes, 4 sessions per week). Only one randomized leg was used for training. Biopsies from the *vastus lateralis* were taken before after the training period. RNA was extracted and sequenced from this tissue as described in ^19^. The resulting count data were uploaded to, and are publicly available in, the Gene Expression Omnibus under the series GSE60590.

Intraindividual changes in gene expression were assessed using the approach described in the section “Differential Analysis” above. Briefly, the gender (male or female) and timepoint (pre- or post-exercise) variables were first concatenated and used in an intercept-free model that also included dominant leg status (i.e. was the randomly chosen leg the subject’s dominant leg) and BMI as additional factors to control for. Subsequent contrasts were fit to assess changes in gene expression as a result of exercise within the male and female subgroups. Fitting was performed using *edgeR::voomLmFit*, blocking on subject ID to account for intraindividual correlation. Gene set testing with cameraPR was performed as described in the section “Gene Set Testing” above. Gene set-level information for GN at the 8-week time-point was used in the comparisons between humans and rats.

### Timewise Trajectory Analysis

#### Fuzzy C-means analysis

Fuzzy C-Means clustering was performed separately by sex after combining the gastrocnemius proteomics, transcriptomics, and metabolomics DA results. Specifically, a matrix of z-statistics from the trained vs. SED comparisons for a particular sex was used as the input. Genes or RefMet metabolite IDs formed the rows of the matrices, while the trained vs. SED comparisons formed the columns. For each sex, only those features with no missing values in any of the 4 z-score columns were retained. A column was included for the SED group and initialized to 0. Since nearly all features were used for clustering, a total of 12 clusters were chosen for both sexes. A higher number of clusters better separates features that do not respond to ExT from those that do, resulting in the discovery of at least several clusters with distinct trajectories in both sexes.

To determine the biological relevance of each cluster, we applied the non-parametric version of CAMERA-PR to test for localization of gene sets or RefMet chemical subclasses. Upper-tail tests were performed separately for each of the 6 combinations of ome and sex with TMSig::cameraPR.matrix using non-default parameter values use.ranks=TRUE and alternative=”greater”. Matrices of cluster membership scores with clusters as columns and features (genes or RefMet metabolites) as rows were used as the input, along with the molecular signatures described previously. The membership scores are probabilities of cluster membership, with the scores across clusters for each feature summing to unity. For each sex and ome, p-values were adjusted across all molecular signatures and clusters using the BH method to control the FDR. FDR < 0.05 was used to determine which molecular signatures were significantly localized to each cluster.

Bubble heatmaps were generated for each combination of ome and sex to display the top molecular signatures localizing to each cluster. If no terms significantly localized to a cluster, the top 2 terms with adjusted p-values in [0.05, 0.1) were included.

## Data Availability

Processed data and analysis results are available in the *MotrpacRatTraining6moMuscleData* R package (https://github.com/MoTrPAC/MotrpacRatTraining6moMuscleData). MoTrPAC data is publicly available via motrpac-data.org/data-access. Data access inquiries should be sent to. Additional resources can be found at motrpac.org and motrpac-data.org. The human RNA-seq count data from the EpiTrain study can be publicly accessed at https://www.ncbi.nlm.nih.gov/geo/query/acc.cgi?acc=GSE60590.

## Code Availability

R scripts detailing all processing and analysis steps are contained within the *data-raw/* folder or in the vignettes and articles of the *MotrpacRatTraining6moMuscleData* R package (https://github.com/MoTrPAC/MotrpacRatTraining6moMuscleData). For a list of the core R/Bioconductor packages used, please consult the Software section of the Supplementary Methods.

## Ethical Statements

The authors declare no conflicts of interest. All animal experiments were approved by the Institutional Animal Care and Use Committee (IACUC) at the University of Iowa Carver College of Medicine and were performed in compliance with institutional and federal ethical guidelines for animal research (NIH Guide for the Care and Use of Laboratory Animals).

## Acknowledgements

The MoTrPAC Study is supported by NIH grants U24OD026629 (Bioinformatics Center), U24DK112349, U24DK112342, U24DK112340, U24DK112341, U24DK112326, U24DK112331, U24DK112348 (Chemical Analysis Sites), U01AR071133, U01AR071130, U01AR071124, U01AR071128, U01AR071150, U01AR071160, U01AR071158 (Clinical Centers), U24AR071113 (Consortium Coordinating Center), U01AG055133, U01AG055137, U01AG055135, U01AG070959, U01AG070960, and U01AG070928 (Pre-Clinical Animal Sites).

---

**Figure S1.**
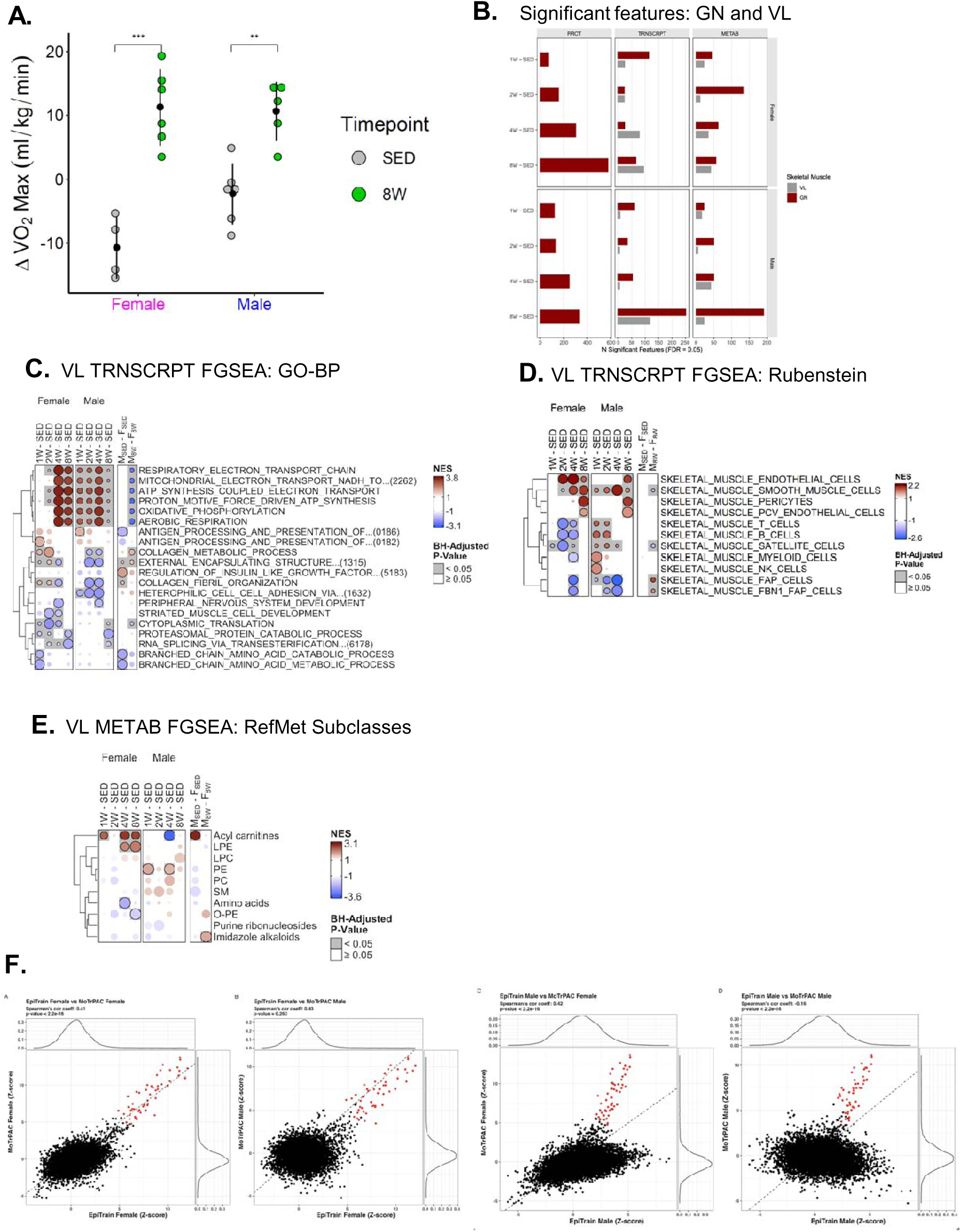
(A) Representative dotplots of changes in VO_2_max (mL/kg/min) over 8-weeks in 8W and SED rats. All changes are represented as change from baseline measurements (mL/kg/min). Data is grouped by sex and color coded by timepoint (SED or 8W). Brackets represent t-test p < 0.01 ** or p < 0.001***. (B) Bar plot representing the number of differential (DA) proteins (PROT), transcripts (TRNSCRPT), and metabolites (METAB) by sex in 1W, 2W, 4W, or 8W groups relative to SED. Here, DA features (BH-adjusted p-value <0.05) are colored as red bars in the GN and gray bars in the VL. (C) Heatmap representing timewise change in Gene Ontology (GO) Biological Processes (BP) of transcriptomics data by contrast in the VL muscle performed using FGSEA. (D) Heatmap of timewise changes of FGSEA gene set enrichments in the VL transcriptomics data curated using a reference gene set from skeletal muscle single cell transcriptomics data from Rubenstein et al. ^13^. (E) Metabolomics pathway enrichments in the VL grouped according to RefMet class analysed by FGSEA. (F) Comparison of transcriptomic responses to exercise in humans and rats at the pathway level. Comparison of cameraPR results for (far left) EpiTrain female versus MoTrPAC female rats; (middle left) EpiTrain female versus MoTrPAC male rats; (middle right) EpiTrain male versus MoTrPAC female rats; (right) EpiTrain male versus MoTrPAC male rats. The dashed line represents y = x. Points highlighted in red indicate gene sets with the highest average Z-score across all comparisons (see text for details). In all bubble heatmaps (panels C-E), a black outline and gray shading indicates statistical significance (BH-adjusted p-values <0.05). In the sex-specific timewise contrasts, red indicates positive enrichment and blue negative enrichment, whereas in the M–F contrasts, blue indicates enriched in females and red indicates enriched in males.

**Figure S2.**
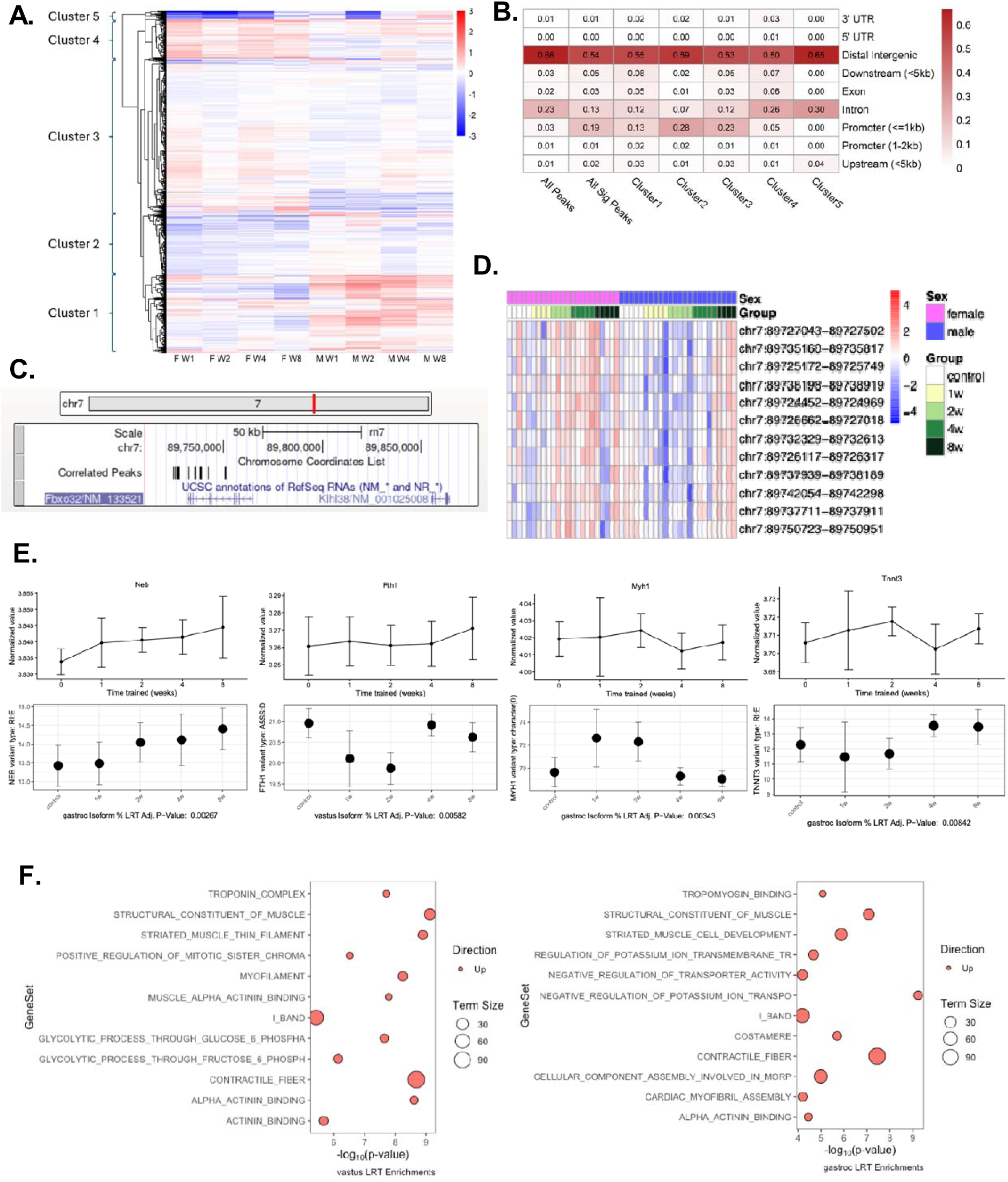
(A) Heatmap of DAR female and male Log2FC responses to exercise training. DARs are hierarchically clustered into five clusters representing distinct patterns of DAR training responses. (B) Gene-body region annotation fractions for all peaks, all DARs, and the DARs in each cluster (shown in Figure 2A). (C) Rn7 gene track for the region of chr7 containing Fbxo32, Klhl38 and the set of correlated peaks from LV10. (D) Heatmap of z-scored accessibility patterns across all muscle samples for top peaks associated with PLIER LV10. (E) Normalized RNAseq transcript expression (top) for alternatively spliced transcript isoforms (bottom); a select set of genes with adjusted p-value <0.05 from the likelihood ratio test (LRT) is presented. The transcript isoform percent-spliced-in is described on the y-axis, indicating what proportion of all isoforms for the transcript the alternatively spliced isoform represents. The type of alternative splicing event is also described (RI: retained introns; SE: skipped exons, AFE: alternative first exons; A5SS/A3SS: alternative 5’ or 3’ splice sites; character(0): Other). (F) Enrichment analysis of differential splicing activity in transcriptomic data via CAMERA-PR. For each gene, the transcript isoform with the greatest test statistic was chosen.

**Figure S3.**
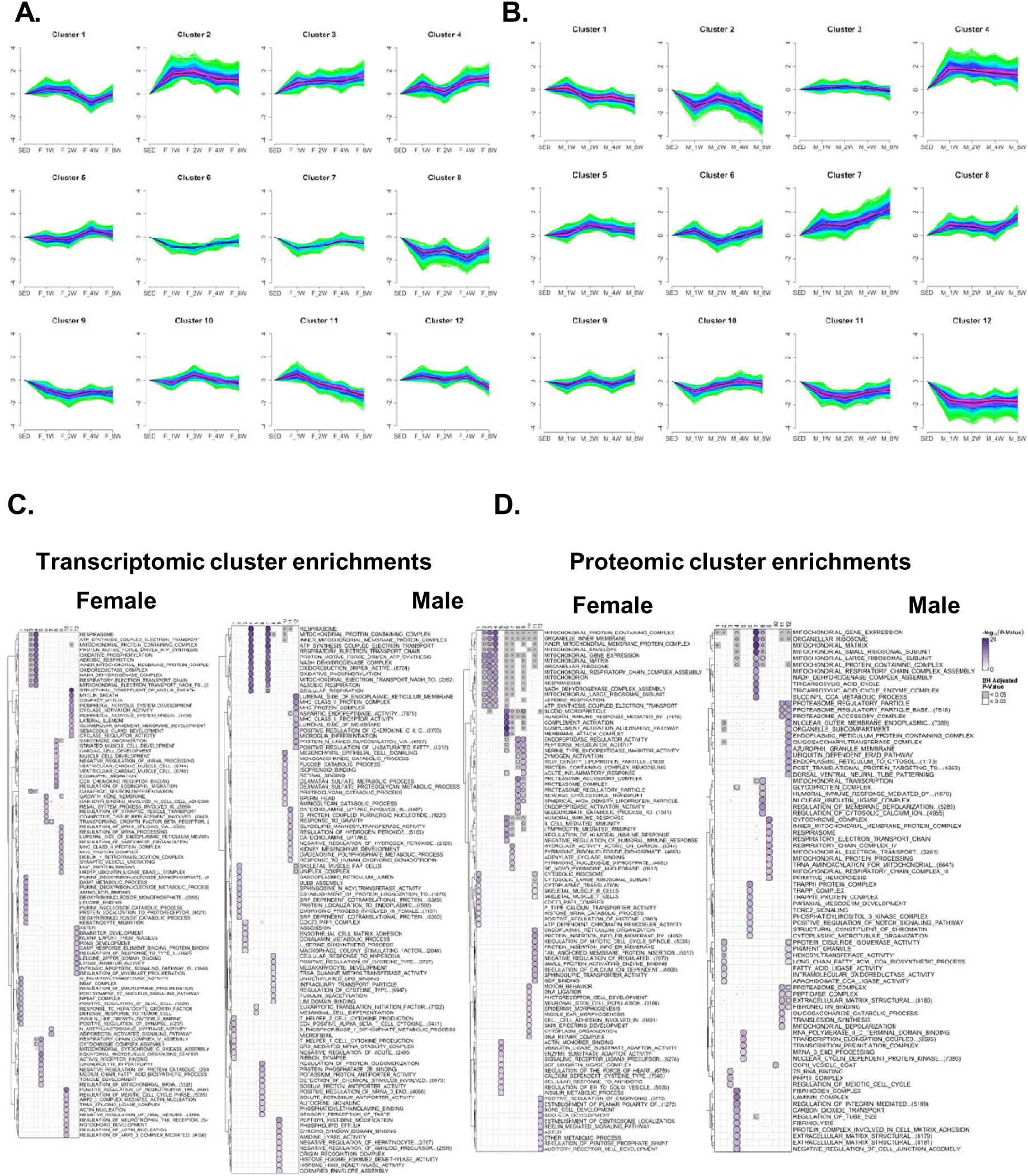
Fuzzy C-means trajectories in females (A) and males (B) for GN metabolomics (METB), transcriptomics (TRNSCRPT), and proteomics (PROT) across training timepoints. Where the y-axis indicates magnitude of features across timepoints, where the dark shading indicates the distribution centroid. (C) Transcriptomic ORA GO enrichment analysis for features localizing to female and male clusters corresponding to Figures S3A and S3B. Gray shading indicates adjusted p-value <0.05. (D) Proteomic ORA GO enrichment analysis for features localizing to female and male clusters corresponding to Figures S3A and S3B. Gray shading indicates adjusted p-value <0.05.

**Figure S4.**
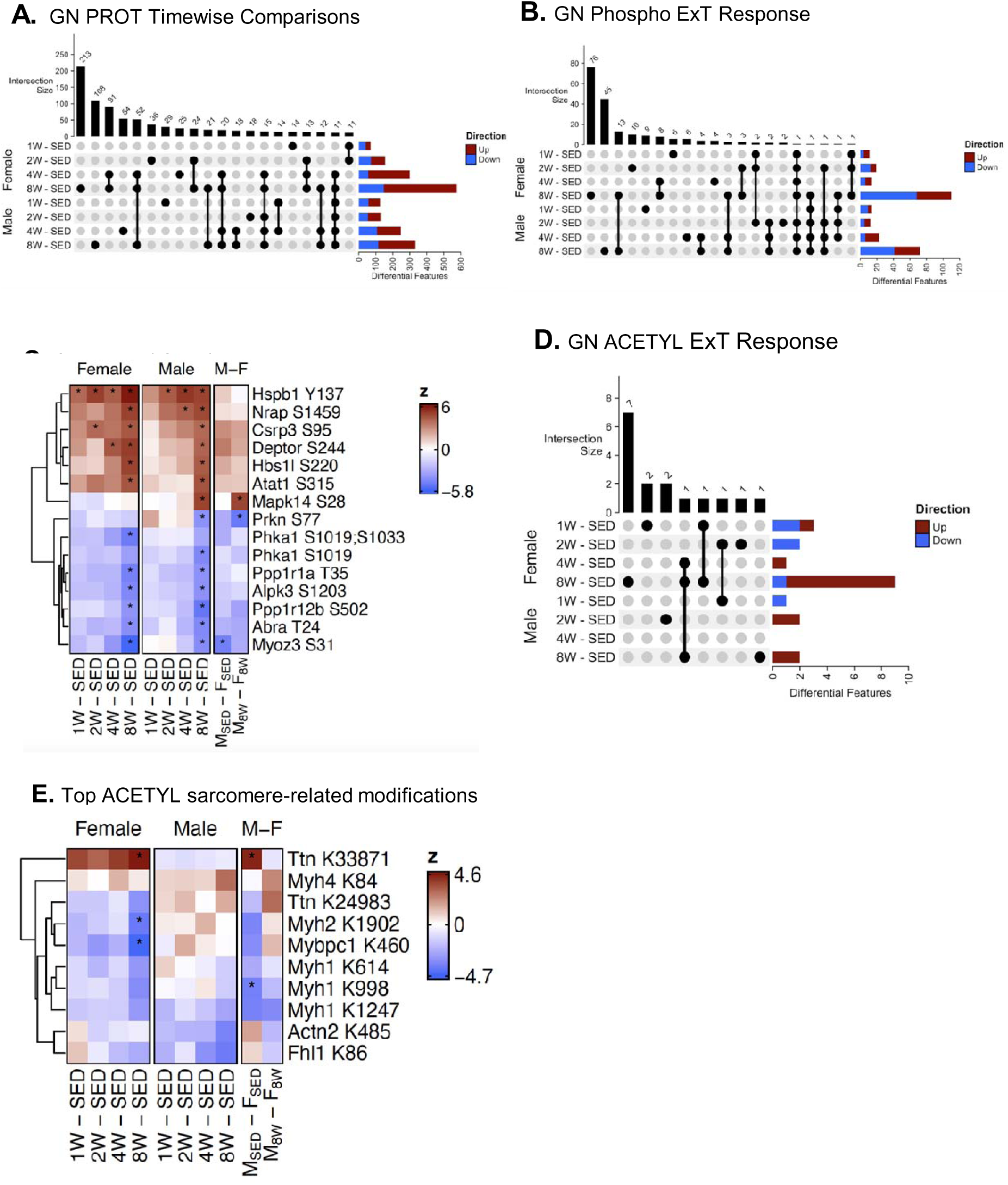
(A) UpSet plot displaying the number of significantly different (BH-adjusted p-value <0.05) proteomic features per training timepoint in female and/or male rats. A line indicates shared DA features across contrasts in males and females, where the number of shared features is indicated as intersection size (top) and the number of DA features per timepoint and sex is indicated on the right. (B) UpSet plot of DA phosphoproteomic features as described in panel A. (C) Feature-level heatmap of select differential phosphoproteins based on biological relevance; * indicates significance at BH-adjusted p-value <0.05 per contrast. (D) UpSet plot of DA acetylome features as described in panel A. (E) Feature-level heatmap of acetylation modifications detected across select sarcomeric proteins; * indicates significance at BH-adjusted p-value <0.05.

**Figure S5.**
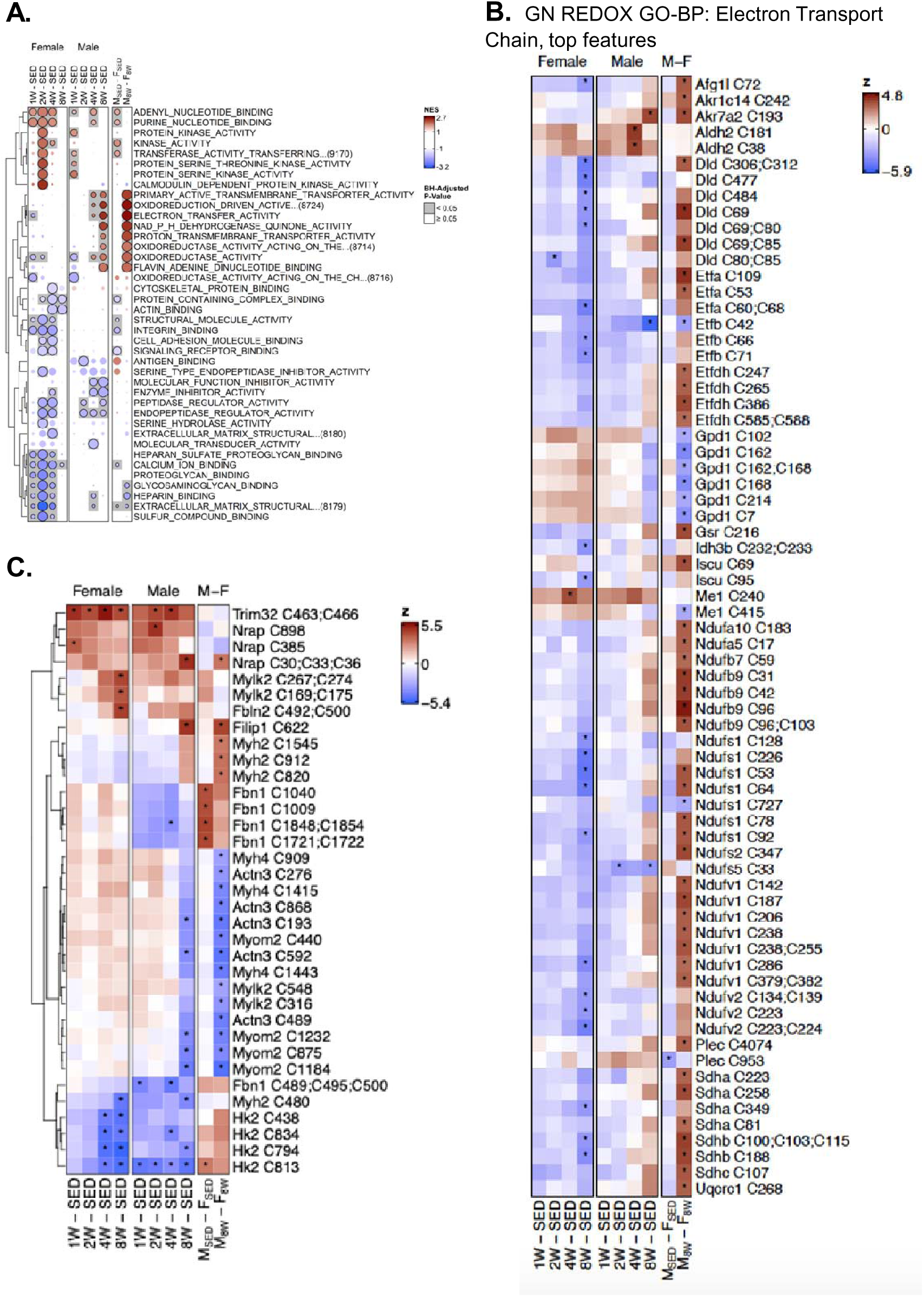
(A) Heatmap displaying the top most significant GO-CC pathway enrichments from FGSEA analysis of the GN cysteine oxidation proteomics dataset; used to contrast the manually curated enrichments presented in Figure 5A. (B) Feature-level heatmaps showing top most significant cysteine oxidation proteomics changes to proteins within the GN REDOX GO-BP: Electron Transport Chain pathway displayed in Figure 5A. (C) Feature-level heatmaps showing cysteine oxidation proteome modifications to select proteins of biological relevance; * indicates significance at BH-adjusted p-value <0.05.

**Figure S6.**
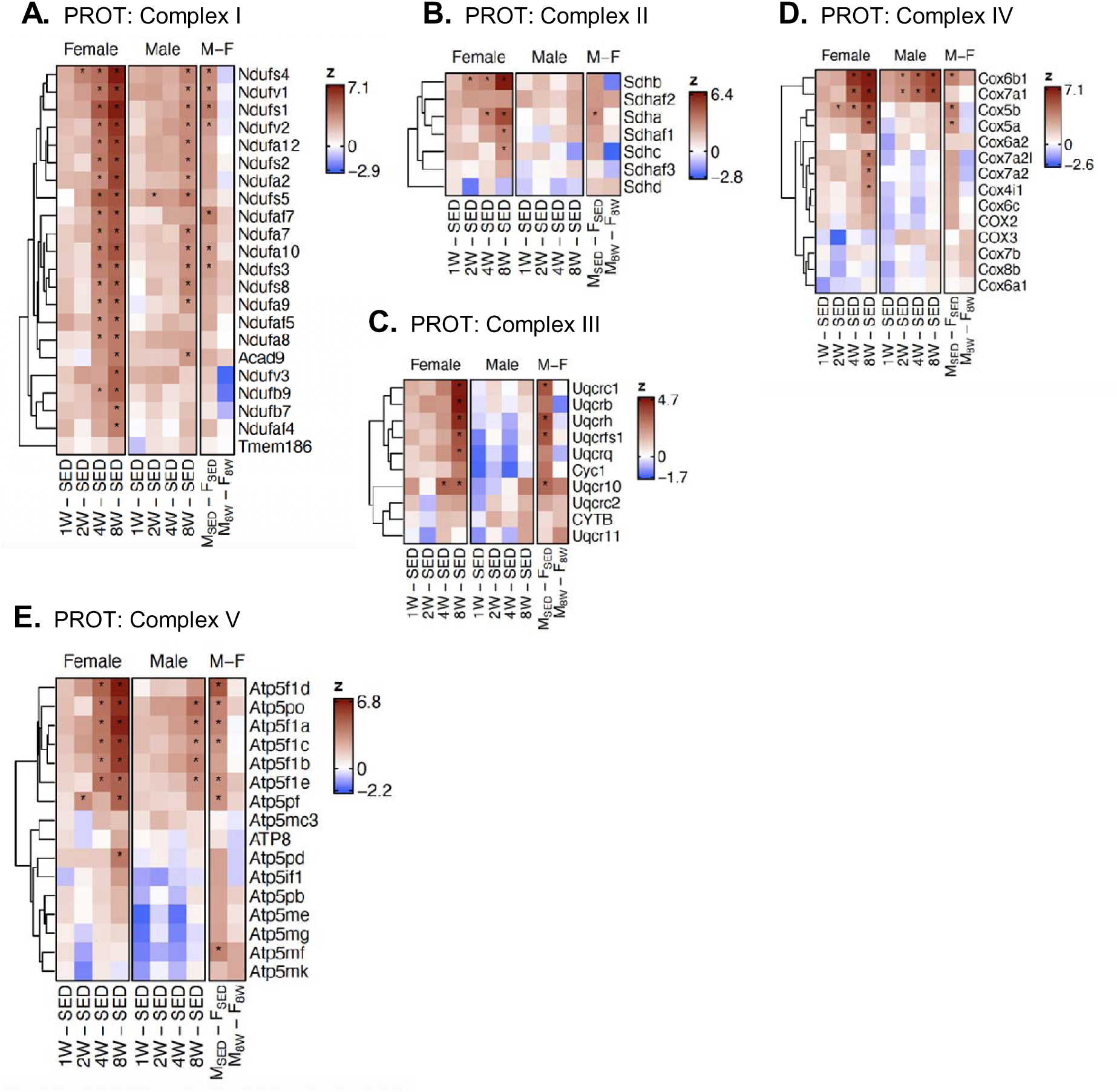
(A-D) Heatmaps displaying the z-score calculated from proteomics differential analysis showing proteins annotated to mitochondrial Complex I (A), Complex II (B), Complex III (C), Complex IV (D) and Complex V (E); * indicates significance at BH-adjusted p-value <0.05 per contrast.

**Figure S7.**
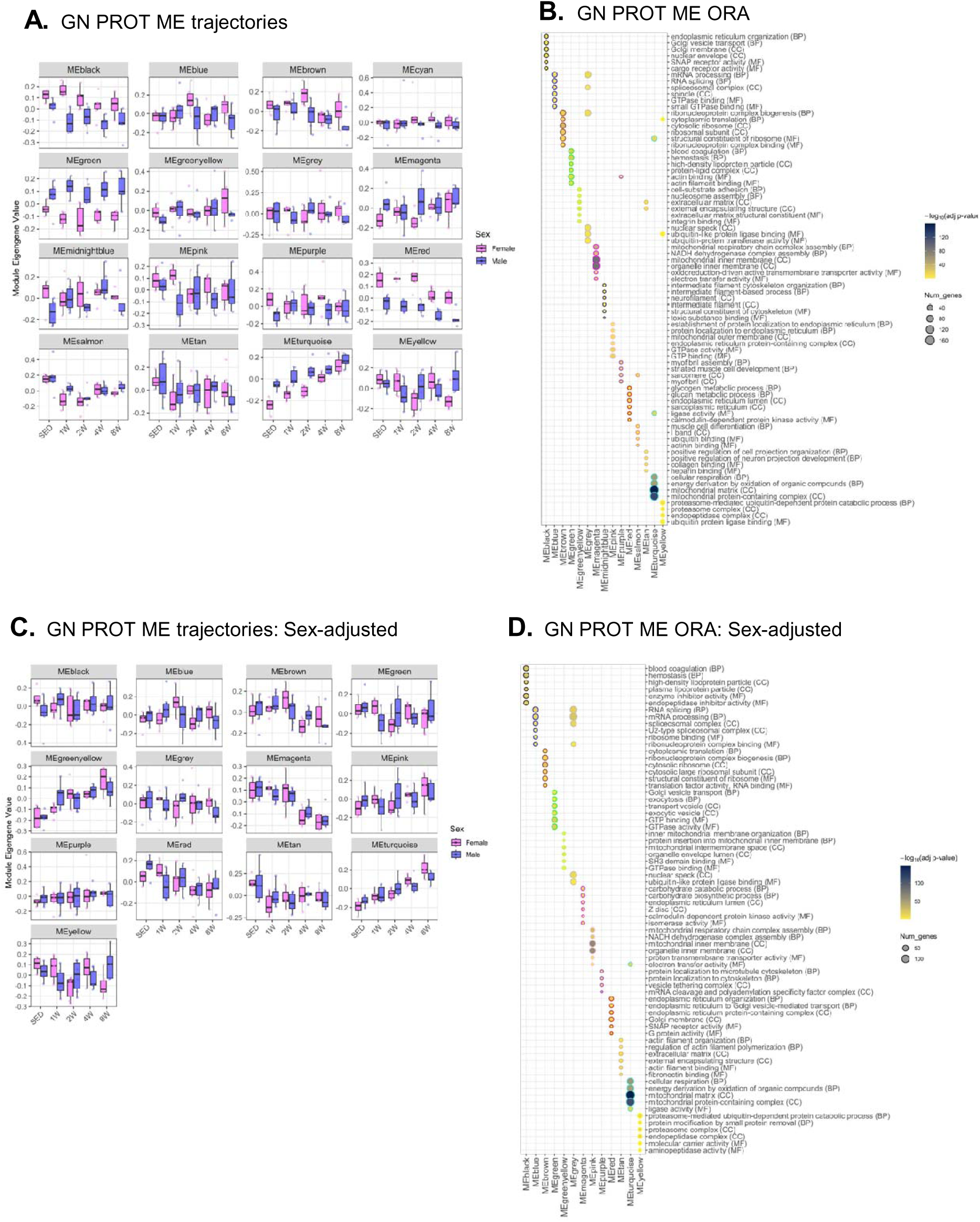
(A) Box plots of ME trajectories from weighted gene co-expression network analysis (WGCNA) in females (pink) and males (blue) without adjusting for sex. (B) Over-representation analysis (ORA) of the proteins contained in all WGCNA MEs generated without adjustment for sex. (C) Box plots of ME trajectories from weighted gene co-expression network analysis (WGCNA) in females (pink) and males (blue) with adjustment for sex. (D) Over-representation analysis (ORA) of the proteins contained in all WGCNA MEs generated with adjustment for sex.

## Notes

https://motrpac-data.org/

### Competing Interest Statement

The authors have declared no competing interest.

